# When the beat drops - The functional anatomy of cardiac-induced sensory attenuation of auditory processing

**DOI:** 10.64898/2026.03.24.713922

**Authors:** Andrew D Levy, Peter Zeidman, Karl Friston

## Abstract

Sensory processing is continuously shaped by internal bodily states, yet the neural mechanisms underlying this interoceptive-exteroceptive integration remain poorly understood. Predictive processing theories propose that bodily states modulate perceptual inference through precision-weighting (the contextual adjustment of prediction error gain according to sensory reliability), but empirical validation with neurobiologically realistic models has been lacking. We address this gap by combining cardiac phase-locked magnetoencephalography with systematic dynamic causal modelling to test competing mechanistic hypotheses about systolic-induced sensory attenuation. Using an auditory oddball paradigm, we observed selective suppression of deviant responses during cardiac systole (200-250ms post-stimulus), affecting prediction errors from unexpected tones whilst sparing expected tones. To identify the underlying synaptic mechanisms, we implemented three methodological innovations: (1) systematic comparison across 20 architectures representing precision-weighting (via intrinsic gains and/or modulatory connections), sensory gating (via forward connections), and predictive suppression (via backward connections) hypotheses; (2) Bayesian model reduction testing all 256 parameter configurations within the winning architecture to handle distributed model evidence; and (3) sensitivity analysis quantifying both direct effects and second-order interactions across the cortical hierarchy. Model comparison decisively favoured precision-weighting implementations (>99.99% posterior probability), with Bayesian model averaging revealing distributed gain control: superficial pyramidal self-inhibition in primary auditory cortex (94%) and inferior frontal gyrus (100%), inhibitory interneuron modulation in superior temporal gyrus (99%), and top-down modulatory connections from superior temporal to primary auditory cortex (96%). Critically, sensitivity analysis demonstrated that intrinsic inhibitory mechanisms exerted order-of-magnitude larger effects than hierarchical modulatory connections, with superior temporal gyrus emerging as an integration nexus showing extensive parameter interactions. These findings provide the first empirical validation of precision-weighting mechanisms during cardiac-sensory integration, establishing that systolic attenuation operates primarily through coordinated local inhibitory gain control rather than hierarchical (expected) attentional modulation. This modelling framework bridges computational theories of interoceptive-exteroceptive integration with laminar-specific cortical mechanisms, offering a generalisable methodology for testing predictive coding hypotheses about embodied perception.

## 1 Introduction

The mind and body are deeply interwoven in shaping sensory experiences, with internal physiology and external stimuli continuously interacting to influence perception and behaviour. This confluence can be obvious, as when internal drives like hunger or thirst compel behaviours to restore homeostasis (Strigo and Craig 2016), or more covert, where visceral states affect sensory sensitivity (Sobel et al. 1998), active sampling (Galvez-Pol, McConnell, and Kilner 2020; Kunzendorf et al. 2019), and motor planning and execution (Galvez-Pol et al. 2022). Visceral influences on perception and cognition can be either facilitating (Garfinkel et al. 2021) or inhibiting (Gianaros and Wager 2015). What is perhaps most striking is that despite the ongoing activity of our internal physiology and its measurable effects on neural processing, how little it features in our conscious awareness.

Here, we focus on one such interaction, the influence of the cardiac cycle on auditory processing, cast in terms of predictive processing. The cardiac cycle comprises two phases: systole and diastole. During systole, ventricular contraction ejects blood into the aorta, creating a transient wave-like increase in blood volume that travels throughout the body. During diastole, the ventricles relax and refill as tissue blood volume returns to baseline. Perceptual sensitivity is often reduced during systole (Al et al. 2020; Louisa Edwards et al. 2001; L. Edwards et al. 2009; Pramme et al. 2016), but see (Garfinkel et al. 2021). The influence of the cardiac system on exteroceptive processing has traditionally been attributed to increased baroreceptor activity inducing cortical inhibition (Bonvallet, Dell, and Hiebel 1954; Rau et al. 1993). However, given the extensive cardiovascular-nervous system interactions, multiple processes likely contribute to cardiac phase-dependent modulation of sensorimotor processing. One can take a broader integrative perspective wherein rapid physiological changes during systole introduce distortionary effects on sensory transduction, transmission, and processing through multiple parallel mechanisms. These interactions can be coarsely categorised into three types: cardio-ballistic interactions arising from pulsatile forces that cause tissue displacement (Thompson and Malina 1959), neuron firing (Uddin 2020), and altered motor control (Sosnoff et al. 2011); cardio-phonic effects from acoustic signals generated by cardiac activity near the ear (Soderquist, Lindsey, and Harter 1973); and neurovascular coupling involving the gain control of neurons as a function of metabolite levels (Lauritzen 2001; Moore and Cao 2008).

This perspective casts cardiac-neural interactions in signal-processing terms where sensory stimuli constitute a signal, whilst systolic-induced perturbations introduce structured noise that degrades sensory processing. Understanding how perception remains stable despite this periodic noise is naturally addressed within a predictive processing framework. Here, perception emerges from hierarchical Bayesian inference whereby higher cortical levels predict lower-level sensory representations, and deviations from these predictions generate prediction errors that ascend the hierarchy (K. Friston 2005; Rao and Ballard 1999). Through iterative belief updating, prediction errors refine internal models to minimise future discrepancies between predictions and sensory input.

The optimality of belief updating hinges on appropriately weighting prediction errors according to their reliability. In predictive coding, this is formalised as precision weighting: a measure that quantifies the certainty associated with a prediction error and acts as a gain control on ascending error signals (Feldman and Friston 2010; Parr and Friston 2019). The assignment of precision involves both predictive and estimative mechanisms. At a prospective level, precision assignment reflects prior expectations about which sensory channels will provide reliable information, thereby determining the allocation of attentional resources to different inputs (K. Friston and Kiebel 2009). At a retrospective level, precision can be estimated from the statistical properties of prediction errors as they are generated, such as their variance or temporal consistency, enabling the system to discount signals that prove unreliable in the moment (K. Friston 2005; Parr and Friston 2019).

This process of selectively enhancing inputs expected to provide the highest information gain is effectively synonymous with selective attention. Or as we pursue here, the complementary process of sensory attenuation. Where attenuation is a result of down-weighting sensory channels through reduced precision assignment. Such a mechanism provides a natural computational account for systolic-induced sensory attenuation. Specifically, neuronal noise induced during systole would result in prediction errors generated during this period being assigned lower precision (relative to diastole), leading to attenuated belief updating (Allen et al. 2022). This would produce a momentary “attending-away” during systole analogous to the attenuation that occurs during saccadic eye movements (Perrinet, Adams, and Friston 2014). This account is consistent with proposals that cardiac-induced sensory attenuation reflects the conservation of limited attentional resources (Berntson and Khalsa 2021; Critchley and Garfinkel 2018; Herman et al. 2021), but extends such proposals by specifying neurobiologically plausible computational mechanisms instantiated in models of predictive coding (Bastos et al. 2012; K. Friston and Kiebel 2009). Importantly, this precision-weighting framework is not competing with the baroreceptor hypothesis but is inclusive of it, as baroreceptor afferent signals would constitute one source of evidence informing the brain about cardiac state and the reliability of sensory channels during systole.

Allen and colleagues (2022) recently provided a computational active inference account demonstrating how cardiac state could modulate precision estimates and thereby influence sensory processing. However, this theoretical proposal has not yet been empirically validated with neural data. In this study, we sought to establish an initial empirical connection between the theoretical model of predictive coding for systole-induced sensory attenuation and the plausible synaptic and hierarchical neuronal circuit mechanisms underlying precision control. To this end, we implemented a human magnetoencephalography (MEG) study using an auditory oddball paradigm and employed dynamic causal modelling (DCM) to interrogate the neuronal mechanisms of cardiac-phase-dependent sensory processing. Following methodologies from previous attention DCM studies (Auksztulewicz and Friston 2015; Brown and Friston 2013), we examined how systole modulates key parameters implicated in cortical gain control. Specifically, we tested for cardiac-phase effects on intrinsic connection strengths (the gains of superficial pyramidal cells and inhibitory interneurons within cortical regions), extrinsic feedforward and feedback connections (transmitting sensory signals and predictions between hierarchical levels), and modulatory top-down connections that regulate the gain of superficial pyramidal cells and inhibitory interneurons (the proposed neurobiological substrate through which expected information gain is instantiated). If cardiac-induced sensory attenuation operates through precision-weighting mechanisms as proposed, we would expect to observe systematic modulation of the intrinsic gain parameters (SP and II self-inhibition) and/or top-down modulatory connections that regulate these gains.

## 2 Methods

### 2.1 Participants

Twenty-four healthy volunteers (20 female, mean age 27.45 ± 4.15 years, range 21-39) participated in the study. All had no history of psychiatric disorders and reported normal hearing. All gave written informed consent and were paid for participation. Procedures were conducted in accordance with the Declaration of Helsinki (1991) and approved by the local ethics committee.

### 2.2 Experimental Paradigm

We used an adapted roving auditory oddball paradigm, where a deviant tone (frequency change from the preceding tone) becomes a standard through repeated presentation. Tone frequencies varied from 800-1300 Hz in 100 Hz steps (sampled from a uniform distribution). Each tone lasted 80ms with 20ms rise/fall times and was presented binaurally via headphones.

We implemented two modifications to the classic paradigm (**Figure 1**). First, tones were presented in blocks differing in interstimulus interval (ISI: 500ms, 1000ms, or 2000ms), held constant within each block. Second, deviant probability was modulated within blocks using a non-homogeneous Poisson process, creating temporal clustering of deviants.^1^ These modifications accommodated concurrent fMRI acquisition (not reported) and tested for ISI-dependent nonlinearities. For MEG analyses, data were pooled across ISI conditions.

**Figure 1.**
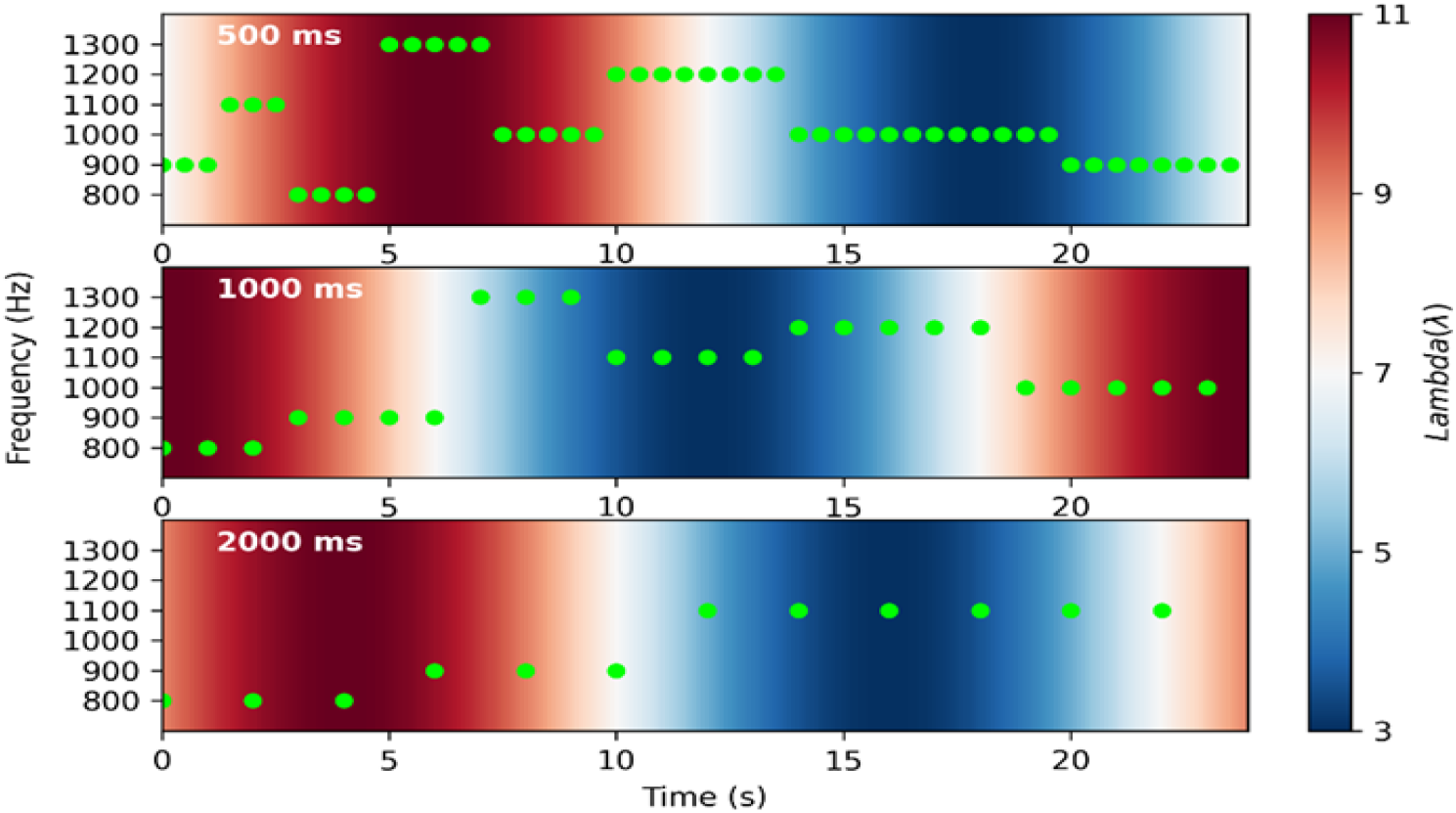
Experimental Paradigm: This figure illustrates the adapted roving oddball design. In this paradigm, a deviant is defined as a change in tone frequency that, through repeated presentations, transitions into a standard tone. The experiment utilised three distinct rates of tone presentation—500ms, 1000ms, and 2000ms—each maintained constant within a single block. Additionally, the frequency of deviant tones was manipulated using a non-homogeneous Poisson process with a sinusoidal modulation, leading to the clustering of deviant tones within a specific segment of the block. The phase of the sinusoid was randomised across blocks, ensuring clustering varied between trials.

Each session comprised nine 32-second blocks with 8-second rest periods. ISI blocks were ordered via Latin square design for balanced exposure. Tone sequences were created offline, and session order was randomised across participants. Each participant completed eight sessions (72 blocks total) over approximately 60 minutes including setup and breaks.

Participants performed an orthogonal cross-hair colour-change detection task to divert attention from tones, pressing a button at each colour change. Changes occurred pseudo-randomly every 6-16s, never coinciding with tone presentation. No participants fell below the 80% hit rate exclusion criterion.

### 2.3 Data Acquisition

MEG data were acquired using a 275-channel whole-head setup with third-order gradiometers (CTF systems) at 1200Hz. Eye movements were recorded via non-ferrous infrared eye-tracking (SR Research). Finger photoplethysmography (PPG) measured the cardiac cycle from the left index finger. Analyses used SPM12 (Wellcome Trust Centre for Neuroimaging, University College London) for MATLAB (Mathworks, Inc.) and custom code for PPG systolic peak extraction and MEG R-peak recovery (Appendix I).

### 2.4 MEG Pre-processing

Raw MEG data were high-pass filtered at 1Hz, downsampled to 150Hz, and low-pass filtered at 48Hz. Auditory event-related fields (ERFs, tone-locked) and cardiac-related fields (CRFs, R-peak-locked during silence) were epoched from -100ms to 400ms and baseline-corrected using the pre-stimulus average. Epochs with Z-scored samples exceeding five standard deviations were excluded.

Cardiac field artefacts (CFA) can contaminate ERFs during systole, confounding cardiac-phase effect interpretations. CFAs were removed using Signal Space Projection (SSP). Spatial components associated with cardiac electrophysiology were identified via Singular Value Decomposition (SVD) of the averaged QRST complex time-windowed to include QRS and T-wave activity (**Figure 2A**). We selected components with eigenvalues exceeding unity (maximum two per window) and excluded them via SSP. Visual inspection confirmed effective CFA representation (average 2.1 components removed). Residual QRST signals post-correction (**Figure 2B**) and auditory ERFs (**Figure 2C-D**) verified no artefact introduction.

**Figure 2.**
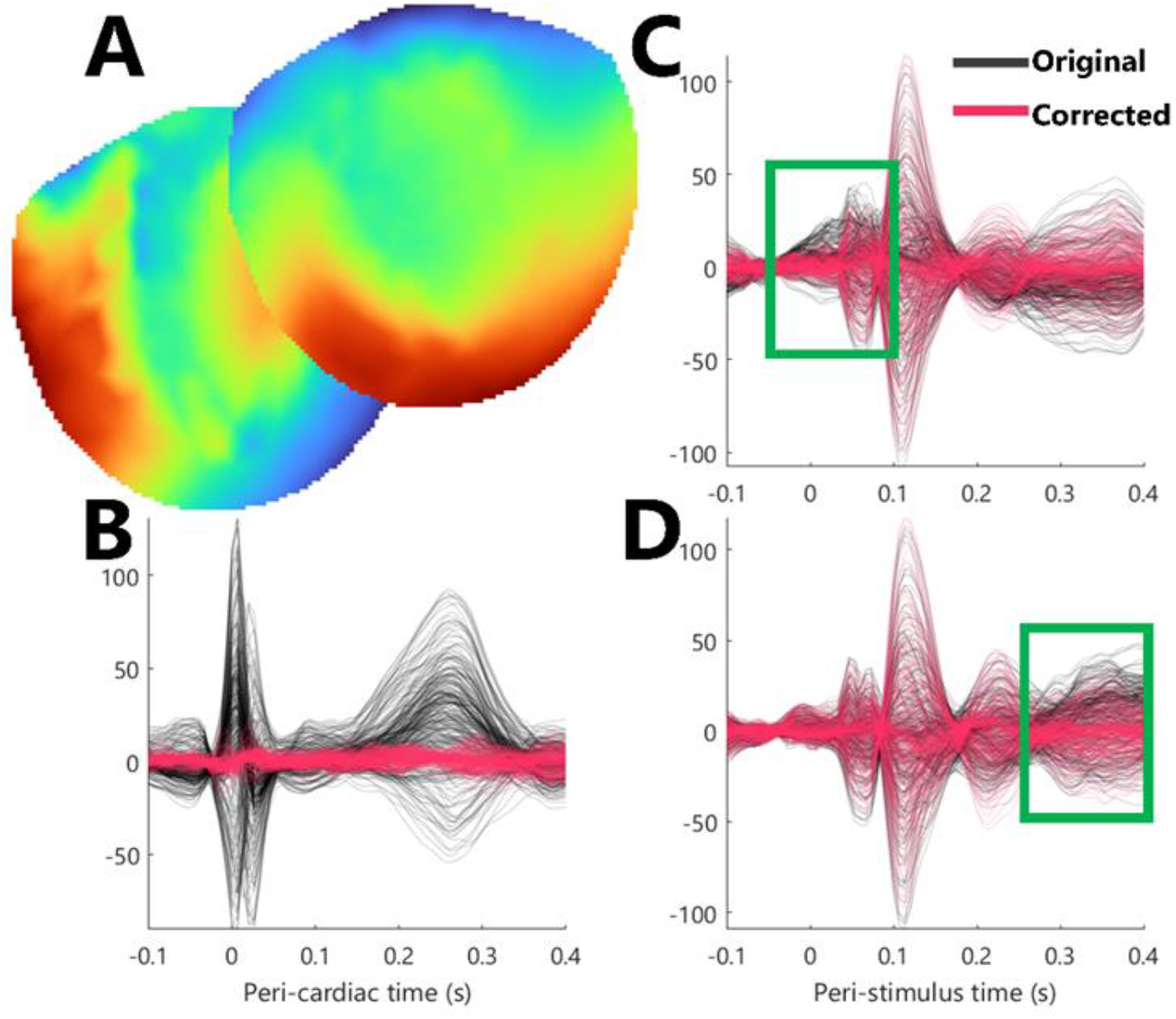
Cardiac Field Artefact Correction: (**A)** Grand averaged spatial topographies of the two principal components associated with the QRST complex. (**B)** Demonstration of the successful removal of QRST complex using signal space projection with responses time-locked to R-peak—responses are the grand average. (**C&D**) Demonstration of the CFA correction on grand-averaged auditory ERFs for tones that occurred during systole (**C**) and tones that occurred during diastole (**D**). Green boxes highlight the specificity of the correction—earlier for systole and later for diastole.

Eye-blink artefacts were corrected similarly, using the vertical eye-tracking channel to identify blinks and removing the primary spatial component via SSP.

### 2.5 Sensor Space Analysis

Cardiac phase was determined by recovering R-peak times from MEG QRST complexes, accounting for pulse wave transit time (PWTT) from finger PPG (Appendix I). Systolic peaks in PPG were identified, average PWTT between MEG R-peaks and PPG peaks calculated, and beat-to-beat R-peaks refined via cross-correlation with subject-specific average QRST complexes. Each cardiac cycle (R-to-R interval) was partitioned into three equidistant segments, excluding 50ms boundaries around R-peaks to avoid contamination. The first partition was systole, the third late-diastole (hereafter diastole). The middle partition (early-diastole) was excluded as it likely contains overlapping processes. Tones were classified by their onset partition. Standards were defined as tones preceded by at least two repetitions of the same frequency. Following artefact rejection and cardiac phase classification, each subject contributed an average of: deviants (systole) 57.2 ± 7.4 trials per condition (range: 39-70), deviants (diastole) 53.3 ± 6.5 trials per condition (range: 42-67), standards (systole) 572.8 ± 27.7 trials per condition (range: 507-616), standards (diastole) 567.2 ± 30.2 trials per condition (range: 476-618).

ERFs were analysed with factors auditory expectation (standards vs deviants) and cardiac phase (systole vs diastole). Subject-averaged ERF time series were converted to 3D images (scalp(x)×scalp(y)×time) and analysed via statistical parametric mapping. Images entered a repeated-measures ANOVA testing main effects and interactions over 2D sensor-space and peristimulus time (100-400ms). Significant effects used random field theory (Kilner, Kiebel, and Friston 2005), testing cluster-size (p<0.001 uncorrected threshold) with family-wise error correction (p<0.05).

### 2.6 Dynamic Causal Modelling

Dynamic Causal Modelling (DCM) for event-related potentials combines biophysically plausible neuronal dynamics with a leadfield forward model for joint source reconstruction and effective connectivity estimation (David et al. 2006). As a Bayesian generative technique, DCM enables systematic model comparison to test hypotheses about underlying neural mechanisms. We applied DCM to test whether systolic-induced attenuation of deviant responses is best explained by precision-weighting.

Testing this hypothesis requires identifying the functional architecture underlying gain control of prediction error-encoding units under visceral modulation. In predictive coding models, superficial pyramidal (SP) cells encode prediction errors (Bastos et al. 2012), and mechanisms modulating SP excitability are key to precision-weighting in vivo (Feldman and Friston 2010; K. Friston 2008; Hertäg and Clopath 2022; Moran et al. 2013). Figure 3 illustrates three candidate mechanisms for precision control: (1) neocortical inhibitory interneuron (II) circuits comprising somatostatin, parvalbumin, and vasoactive intestinal peptide interneurons controlling SP gain (Barron, Auksztulewicz, and Friston 2020; Hertäg and Clopath 2022); (2) extrinsic neuromodulation via acetylcholine, dopamine, and norepinephrine (Alkondon et al. 2000; Froemke 2015); and (3) NMDA receptor-mediated potentiation (Cornford et al. 2019).

**Figure 3.**
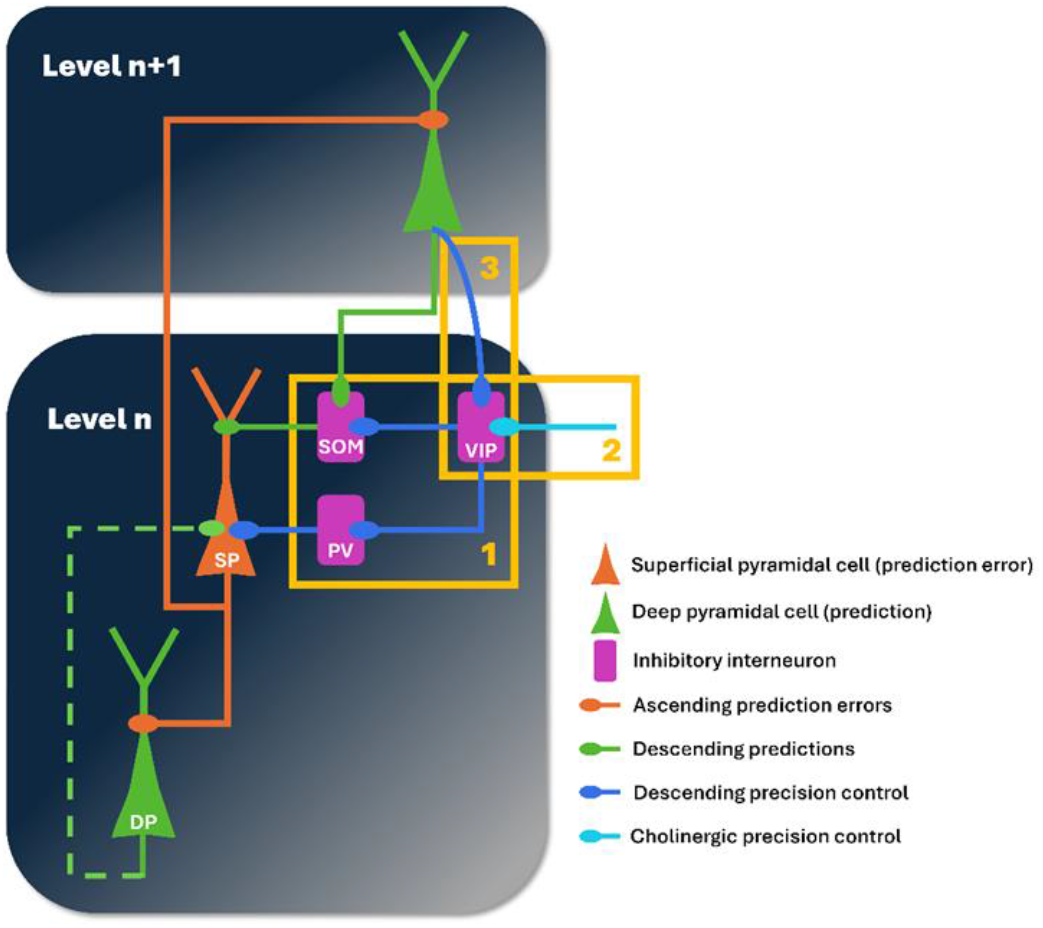
Neocortical microcircuit schematic of descending precision control in predictive coding: (Box1) Intrinsic inhibitory interneuron circuit. (Box 2) Extrinsic cholinergic neuromodulation. (Box 3) Top-down NMDA modulation. Legend: superficial pyramidal (SP), deep pyramidal (DP), somatostatin inhibitory interneuron (SOM), parvalbumin inhibitory interneuron (PV), vasoactive intestinal peptide inhibitory interneuron (VIP)

We conducted systematic model comparison to determine whether systolic-induced attenuation of deviant responses is best explained by precision-weighting of prediction errors (via the gain control mechanisms described above), sensory gating of ascending connections, or enhanced predictive suppression via descending connections.

#### 2.6.1 Sources Specification

Sources comprised bilateral primary auditory cortex (A1), superior temporal gyrus (STG), and inferior frontal gyrus (IFG), the established network underlying mismatch negativity (Garrido et al. 2008; Ishishita et al. 2019; Opitz et al. 2002). Prior locations were informed by multiple sparse priors source reconstruction of grand-averaged responses (1-32 Hz, 200-300ms window) in SPM12 (K. Friston 2008). Based on maximum intensity projections (), MNI coordinates were: rA1 (50,-20,10), lA1 (-50,-20,10), rSTG (60,-15,-10), lSTG (-60,-15,-10), rIFG (49,18,17), lIFG (-49,18,17). Connections followed conventional auditory hierarchy (**Figure 4A**).

**Figure 4.**
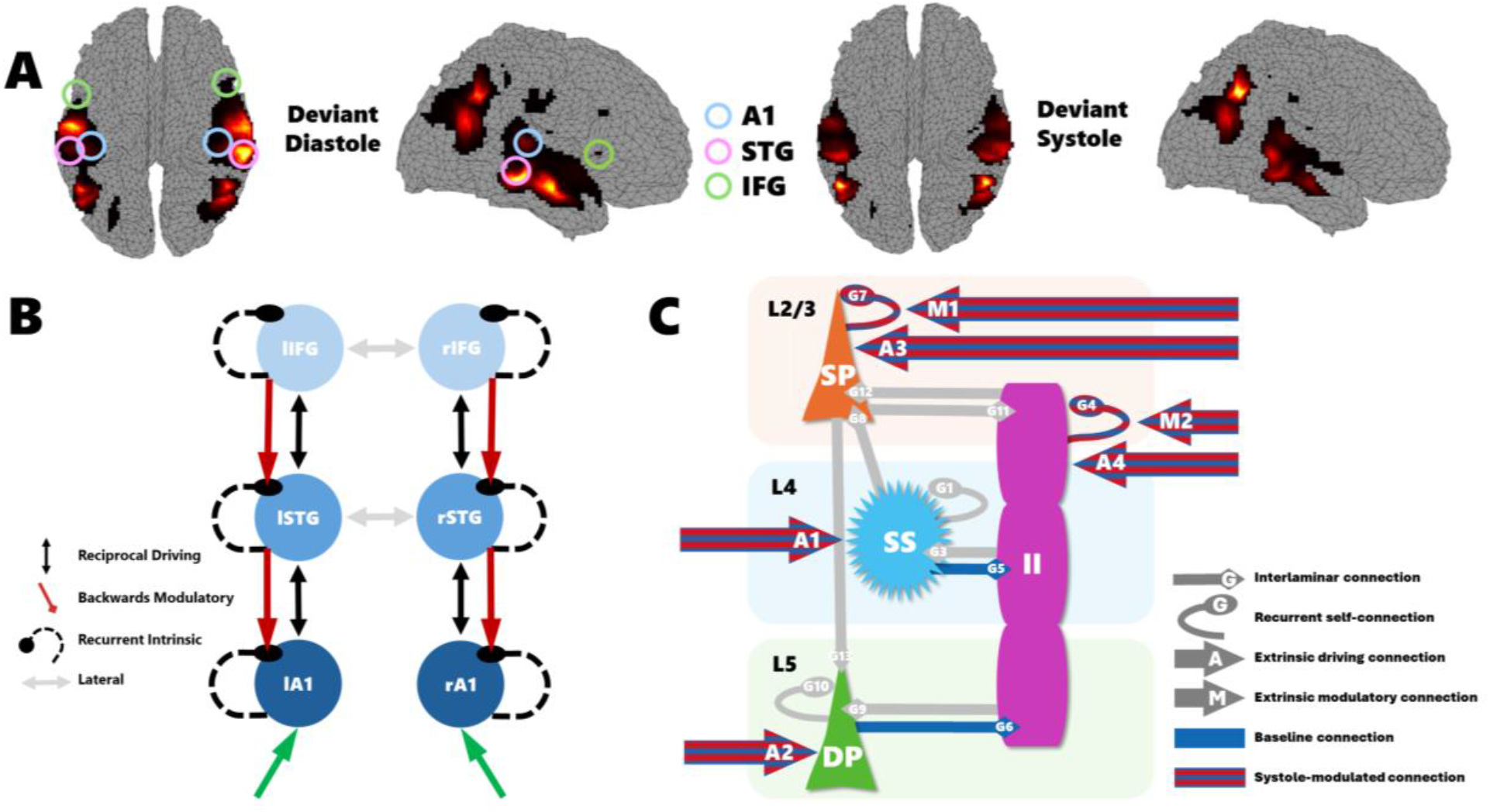
DCM specification: (**A**) Prior source locations identified through multiple sparse prior source reconstruction, (**B**) hierarchical auditory network specification, (**C**) canonical microcircuit model and estimated connections.

#### 2.6.2 Neuronal Model Specification

##### Neural mass model

Each source used the canonical microcircuit (CMC) model, selected for explicitly modelling laminar circuitry implementing predictive coding (Bastos et al. 2012; Shaw et al. 2017). The CMC comprises superficial pyramidal (SP), deep pyramidal (DP), spiny stellate (SS), and inhibitory interneuron (II) populations with established intralaminar and recurrent self-inhibitory connections (Haeusler and Maass 2006; Thomson 2003) (**Figure 4B**). Predictive coding is implemented via asymmetric extrinsic connections: ascending (prediction errors) from SP to SS/DP in higher regions; descending (predictions) from DP to SP/II in lower regions.

##### Intrinsic gain control

II processes are specified via an explicit II population (lumped model of SOM, PV, VIP subtypes with reciprocal connections) and implicit recurrent self-connections. Self-inhibitory connections capture II activity plus other excitability/refractoriness processes (adaptation, spike-rate dependence).

##### Top-down gain modulation

Non-linear modulatory connections allow higher DP cells to modulate lower SP and II self-inhibition (Brown and Friston 2013), capturing slow NMDA receptor modulation of SP gain:

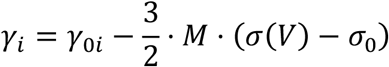

where self-inhibition (*γ*) of recipient SP cells is modulated from baseline (*γ*_0_) by afferent DP firing rates. Firing rates (*σ*(*V*)) are sigmoid functions of depolarisation (*V*); deviations from baseline (*σ*_0_) drive self-inhibition changes. Modulation strength is parametrised by *M* (positive values suppress self-inhibition, increasing gain). We extended this to include II gain modulation, representing top-down control of tonic GABA crucial for selective processing (Auger, Meccia, and Floresco 2017; Bryson et al. 2020; Sanders et al. 2013; Womelsdorf et al. 2014).

##### Model constraints

Moderate symmetry constraints (*r* = 0.6 prior covariances) between hemispheric pairs reduced complexity, focusing on hierarchical rather than hemispheric effects while allowing variations.

#### 2.6.3 Data and Design Specification

DCM analysed grand-averaged ERFs to maximise signal-to-noise, assuming common cardiac-phase mechanisms across participants. Analyses modelled cardiac-specific connectivity changes for deviant responses (0-400ms). Focus on deviants was motivated by sensor-level analyses showing cardiac modulation specific to deviants. Exogenous input was modelled as Gaussian (peak 64ms, SD 16ms) delivered bilaterally to A1. Diastolic onsets served as baseline; systolic onsets as modulatory condition (**Figure 4C**).

#### 2.6.4 Bayesian Model Comparison & Reduction

Posterior estimates depend on model structure (Stephan et al. 2010), requiring Bayesian Model Comparison (BMC) via free energy (model evidence approximation).

##### Bayesian Model Comparison

We conducted systematic model comparison to adjudicate between three candidate mechanisms for cardiac modulation of auditory deviance processing as can be identified within the DCM framework. These were specifically:

***Precision-weighting*** posits that cardiac processes modulate the gain of prediction error units. This predicts cardiac modulation of **intrinsic gain parameters** (I), and/or **modulatory extrinsic connections** (N) where higher cortical areas dynamically adjust lower-level gain through NMDA-mediated modulation.

***Sensory gating*** proposes that cardiac processes such as baroreceptor mediated cortical inhibition attenuate ascending signals. This predicts cardiac modulation of **forward driving connections** (B), reducing bottom-up signal transmission from lower to higher areas.

***Predictive suppression*** suggests cardiac processes enhance descending predictions that ‘explain away’ deviant signals. This predicts cardiac modulation of **backward driving connections** (B), strengthening top-down predictive signals.

We systematically tested all combinations of these three factors [I × B × N]: **intrinsic gains** [I: 1=SP only, 2=SP & II], **driving extrinsic connections** [B: 0=none, 1=forward, 2=backward, 3=both], and **modulatory extrinsic connections** [N: 0=none, 1=DP→SP gain, 2=DP→SP & II gains]. All permutations were considered with the constraint that top-down II modulation (N=2) required intrinsic II gains (I=2), yielding 20 candidate models.

##### Bayesian Model Reduction

The winning BMC model (Model 10: top-down modulatory connections to SP with intrinsic SP and II gains) underwent Bayesian Model Reduction (BMR) (K. J. Friston et al. 2016). BMR tests whether simpler models (parameters fixed to zero) better explain data, effectively pruning to identify essential parameters. We systematically tested all combinations of the following parameters, where each parameter was either included (modulated by systole) or excluded (set to zero) from the model:

- SP gain in A1 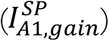
- SP gain in STG 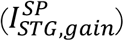
- SP gain in IFG 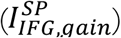
- II gain in A1 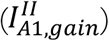
- II gain in STG 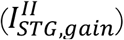
- II gain in IFG 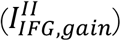
- Top-down modulatory connection from STG to A1 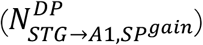
- Top-down modulatory connection from IFG to STG 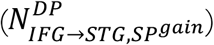

Each parameter represents bilateral connections—inclusion/exclusion applied symmetrically to both hemispheres, consistent with moderate symmetry constraints (*r* = 0.6) and hierarchical focus. Testing all permutations of 8 bilateral parameter sets yielded 2^8^ = 256 candidate models compared via model evidence.

##### Bayesian Model Average

BMR yielded no single winner; posterior probability was distributed across configurations. Bayesian Model Average (BMA), therefore, provided principled inference, averaging parameter estimates weighted by posterior probabilities (Trujillo-Barreto, Aubert-Vázquez, and Valdés-Sosa 2004), accounting for model uncertainty more robustly than single-model conditioning.

##### Sensitivity Analysis

To understand parameter contributions, we performed sensitivity analysis on Model 10; parameter selection was informed by BMR results. First-order and second-order partial derivatives examined how deep pyramidal (DP) and superficial pyramidal (SP) voltages changed with parameter perturbations. We calculated accumulated absolute voltage differentials across the post-stimulus period, characterising sensitivity by magnitude independent of polarity. First-order analysis quantified direct parameter influences; second-order analysis examined interactions, revealing how parameter combinations jointly influence hierarchical responses. We also qualitatively examined peristimulus time courses (SP, SS, II, DP) and reconstructed local field potentials.

## 3 Results

### 3.1 Sensor Space Analysis

We first examined whether cardiac phase modulates auditory processing at the sensor level, focusing on prediction error responses. Event-related fields (ERFs) were analysed using a 2×2 repeated-measures ANOVA with factors auditory expectation (standards vs deviants) and cardiac phase (systole vs diastole). A significant main effect of auditory expectation peaked at 253 ms over centro-temporal channels (peak-level *F*_*max*_ = 83.95; cluster-level *p*_*FWE*_ = 1.57 × 10^−9^; **Figure 5A, Table 1**), consistent with robust mismatch negativity responses. No significant main effect of cardiac phase survived multiple comparison correction (**Figure 5B**), indicating that cardiac modulation does not produce general sensory suppression but rather interacts with expectation-related processing.

**Table 1:** Cluster- and peak-level statistics for significant effects. All peaks survived family-wise error correction (p<0.05). Coordinates are in sensor space (x, y) and peristimulus time (pst)

| cluster |  |  | peak |  |  | x | y | pst |
| --- | --- | --- | --- | --- | --- | --- | --- | --- |
| $p_{FWE\text{ corr}}$ | $K_E$ | $p_{unc}$ | $p_{FWE\text{ corr}}$ | F | $p_{unc}$ | mm | mm | ms |
| Main effect of auditory expectation |  |  |  |  |  |  |  |  |
| 1.57E-09 | 1236 | 3.44E-10 | 1.21E-09 | **83.95** | 1.92E-14 | 47 | -9 | 253 |
| 2.35E-09 | 1202 | 5.14E-10 | 2.98E-09 | **80.49** | 4.72E-14 | -34 | -25 | 260 |
| 3.20E-09 | 1176 | 7.01E-10 | 1.27E-08 | **75.1** | 2.00E-13 | 42 | -14 | 133 |
| 6.35E-08 | 935 | 1.39E-08 | 2.37E-08 | **72.83** | 3.74E-13 | -38 | -25 | 133 |
| 5.28E-07 | 776 | 1.15E-07 | 4.55E-08 | **70.31** | 7.54E-13 | -60 | 8 | 260 |
| 8.65E-07 | 740 | 1.90E-07 | 8.43E-08 | **67.97** | 1.47E-12 | -51 | 29 | 140 |
| 4.62E-04 | 389 | 1.01E-04 | 3.79E-03 | **31.88** | 1.98E-07 | 51 | 40 | 133 |
| 7.57E-04 | 312 | 1.66E-04 | 1.63E-02 | **27.53** | 1.06E-06 | 55 | 45 | 213 |
| 5.81E-03 | 209 | 1.28E-03 | 6.91E-02 | **23.28** | 5.85E-06 | 42 | -19 | 7 |
| Interaction of auditory expectation and cardiac phase |  |  |  |  |  |  |  |  |
| 1.07E-02 | 181 | 2.35E-03 | 1.63E-02 | **27.53** | 1.06E-06 | 38 | -3 | 233 |

**Figure 5.**
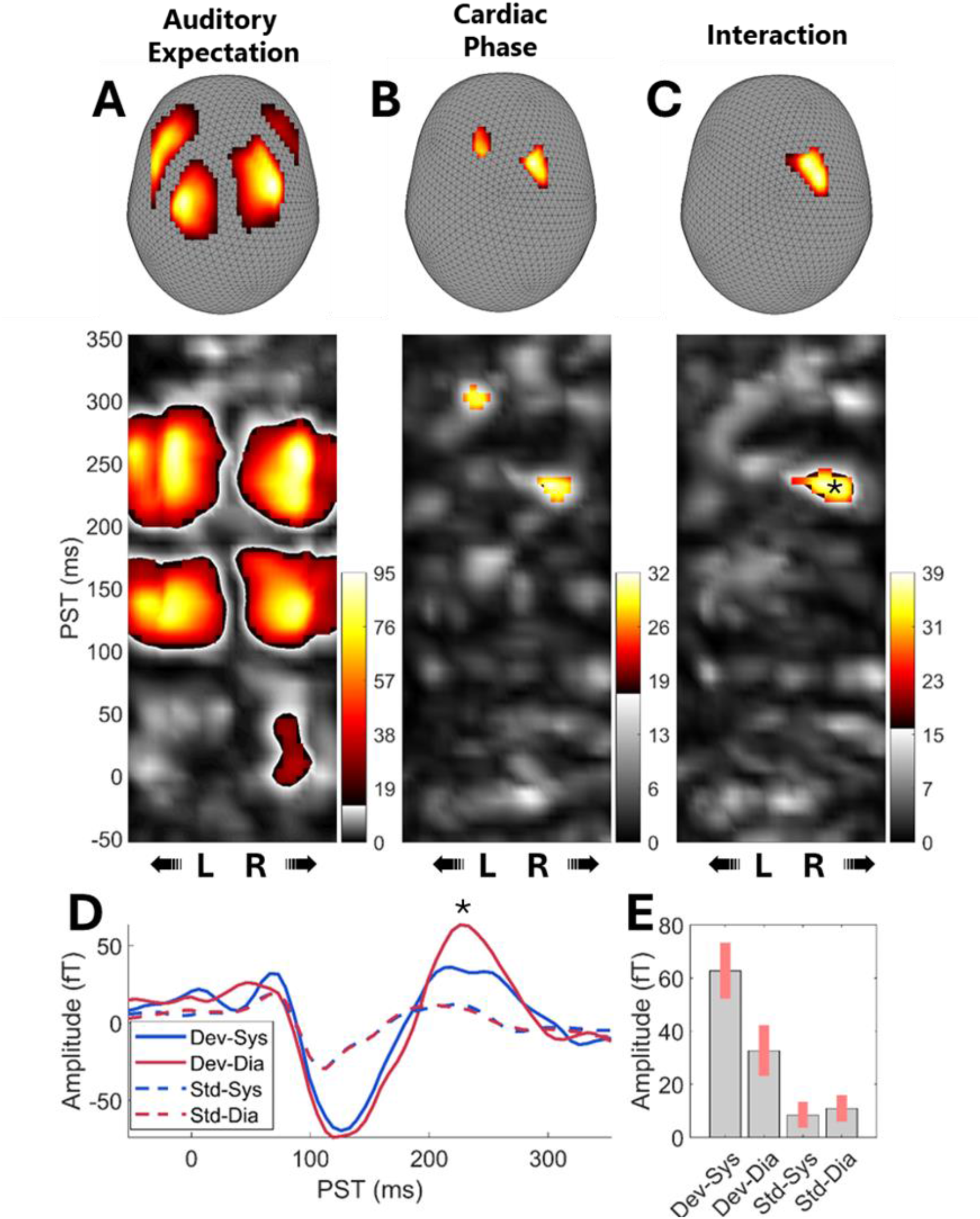
Sensor Space Results: (**A-C**) Maximum intensity projections of thresholded F-contrasts for the (**A**) main effect of auditory expectation, (**B**) main effect of cardiac phase and (**C**) the interaction of auditory expectation and cardiac phase. Peak coordinates survived family-wise error (FWE) correction for multiple comparisons (see Table 1) and are shown at p < 0.001, uncorrected for display purposes. (**D**) ERF at the statistical maxima (38 mm, -3 mm; reconstructed from the GLM coefficients; ‘*’ indicates the peristimulus statistical maxima). (**E**) Coefficient estimates with 90% confidence interval at the statistical maxima (38 mm, -3mm, 233 ms) for the interaction of auditory expectation and cardiac phase.

We identified a significant interaction between auditory expectation and cardiac phase, with peak activation at 233 ms over centro-temporal channels (peak-level *F*_*max*_ = 27.53; cluster-level *p*_*FWE*_ = 0.011; **Figure 5C, Table 1**). This timing corresponds to the late mismatch negativity component associated with prediction error signaling. As shown in **Figure 5D-E**, deviant responses were specifically suppressed during systole relative to diastole. Follow-up paired t-tests examining each tone type separately confirmed that this cardiac phase effect was significant for deviants (peak-level *T*_*max*_ = 5.15; cluster-level *p*_*FWE*_ = 0.016, 38 mm, -3 mm, 233 ms) but not for standards (peak-level *T*_*max*_ = 4.04; cluster-level *p*_*FWE*_ = 0.260). These findings support the hypothesis that systolic-induced sensory attenuation selectively modulates prediction error processing rather than producing general sensory suppression.

### 3.2 Dynamic Causal Modelling

#### 3.2.1 Bayesian Model Comparison

Having established that the cardiac phase specifically modulates deviant responses at the sensor level, we next sought to identify the neural mechanisms underlying this effect. We performed model comparison to adjudicate between three candidate hypotheses about how cardiac phase modulates auditory processing: (1) **precision-weighting** via intrinsic gain parameters and/or top-down modulatory connections, (2) **sensory gating** via modulation of forward driving connections, and (3) **predictive suppression** via modulation of backward driving connections.

We compared twenty candidate models, systematically varying across three factors: **intrinsic gains** (I: SP only vs SP & II), **driving extrinsic connections** (B: none, forwards, backwards, or both), and **modulatory extrinsic connections** (N: none, DP→SP only, or DP→SP & II). These three factors yielded 20 models with the constraint that top-down II modulation (N=2) required intrinsic II gains (I=2). **Figure 6** displays the log model evidence and posterior probabilities for all models.

**Figure 6.**
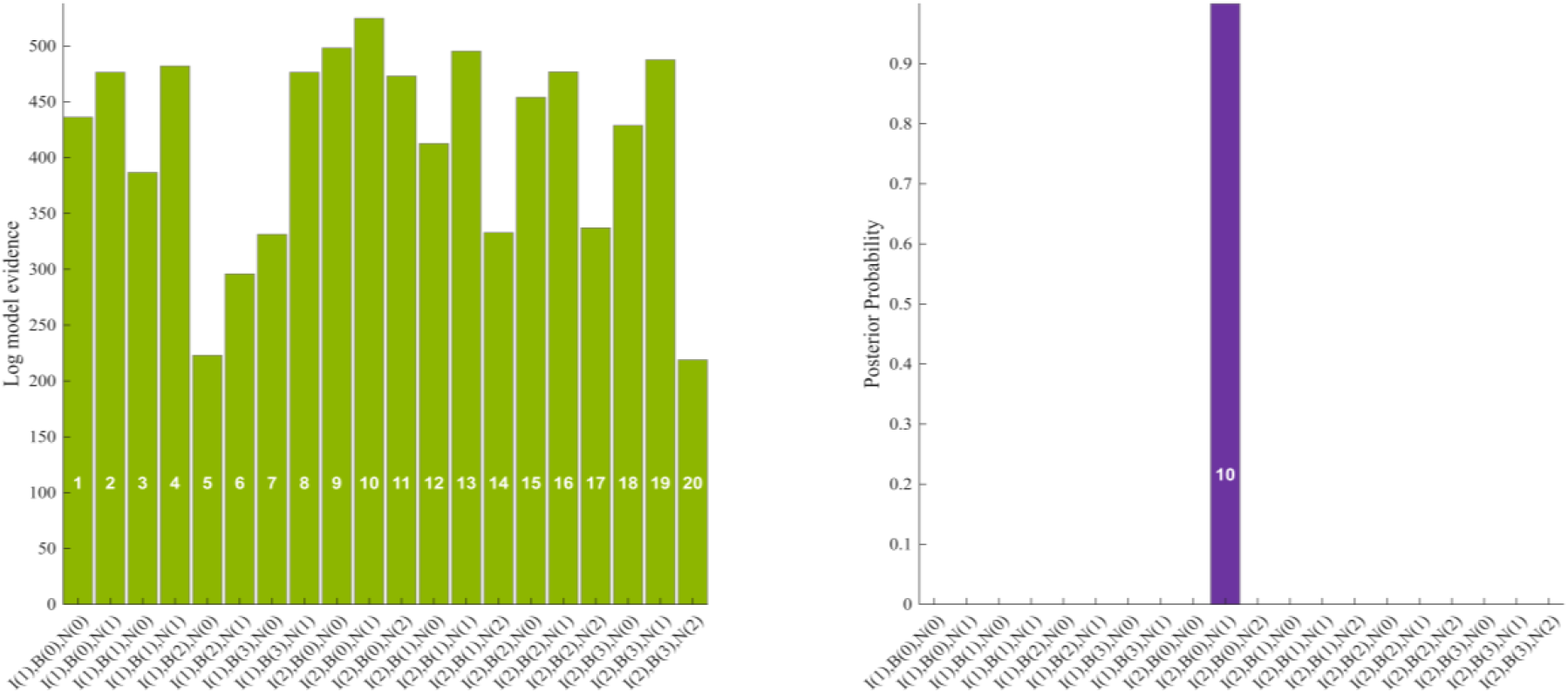
Results of Bayesian Model Comparison of Modulated Connections: Log model evidence was assessed for models with different configurations of cardiac-dependent modulations of extrinsic and intrinsic connections, as follows: **Top-down modulatory connections (N)**: 0 (none), 1 (DP to SP only), 2 (DP to SP & DP to II); **Driving extrinsic connections (B):** 0 (none), 1 (forward), 2 (backward), 3 (both forward and backward); **Intrinsic gains (I):** 1 (SP gain only) or 2 (SP and II gain). The log evidence (left panel) shows that Model 10, with cardiac-dependent top-down modulatory connections targeting only superficial pyramidal cells and incorporating intrinsic gains in both superficial pyramidal and inhibitory interneurons, exhibited the highest posterior probability (right panel).

The analysis revealed strong support for precision-weighting mechanisms. Model 10, incorporating top-down modulatory connections targeting superficial pyramidal cells (N=1) alongside intrinsic gains in both superficial pyramidal and inhibitory interneurons (I=2), achieved the highest model evidence (LME = 524.75) and dominated with a posterior probability exceeding 99.99%. The next-best model (Model 9: LME = 498.25), which included only intrinsic gains without top-down modulatory connections, showed substantially lower evidence (ΔLME = 26.5), indicating decisive support for the inclusion of modulatory connectivity.

Models incorporating driving connections (either forward or backward) showed poor performance. No model with forward driving connections (B=1) achieved meaningful posterior support, decisively rejecting the sensory gating hypothesis. Similarly, models with backward driving connections (B=2) performed poorly, rejecting the predictive suppression hypothesis. The pattern of evidence clearly favoured mechanisms operating through gain modulation rather than through changes in driving connectivity.

These findings indicate that cardiac modulation operates through precision-weighting mechanisms implemented via both intrinsic cortical gains and top-down modulatory pathways. The necessity of including both SP and II intrinsic gains suggests that systolic effects modulate excitability across multiple cell populations within local circuits. The critical addition of top-down modulatory connections (but not driving connections) indicates that cardiac phase involves neuromodulatory gain control rather than directly driving or gating neuronal activity.

#### 3.2.2 Bayesian Model Reduction

We subjected the winning model (Model 10) to Bayesian Model Reduction (BMR) to identify which specific parameters were essential for explaining systolic-induced effects. BMR systematically tested all 256 possible combinations of the 8 bilateral parameter sets (each parameter either included or excluded), comparing their model evidence to determine which subset of parameters best captured the observed data.

BMR yielded no single definitive winning model (**Figure 7A-B**). The highest-ranked model achieved a posterior probability of only 19.8%, with model evidence distributed across multiple competing configurations. However, this distribution was relatively concentrated: the top 12 models (4.7% of model space) accounted for over 90% of the total posterior probability, indicating that whilst no single configuration dominated, support converged on a limited set of parameter combinations. This pattern suggests the underlying mechanism involves contributions from several complementary gain control processes rather than a single critical parameter.

**Figure 7.**
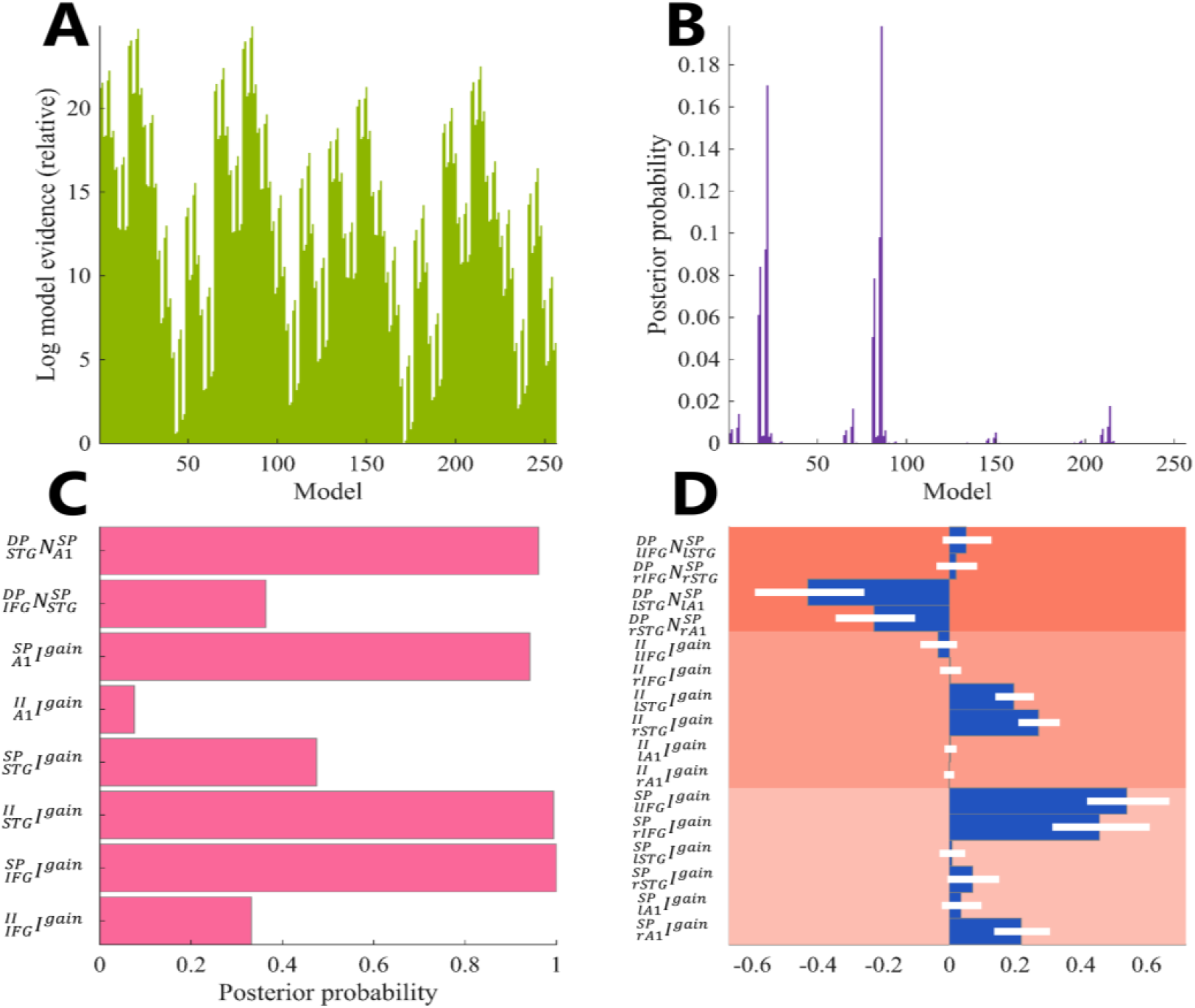
Bayesian Model Reduction and Model Average Results: (**A**) Log model evidence across all 256 candidate models, relative to the lowest-ranked model. (**B**) Posterior probabilities show no single dominant configuration, with the highest-ranked model achieving only 19.8% probability. (**C**) Posterior probabilities of individual parameters from the Bayesian Model Average, indicating confidence that each parameter contributes to systolic-dependent effects. (**D**) Posterior parameter estimates (mean and 95% credible intervals) from the Bayesian Model Average, showing direction and magnitude of systolic-induced modulation for each parameter.

Given the absence of a clear winning model, we based our parameter inferences on the Bayesian Model Average (BMA), which provides a principled approach for handling model uncertainty. The BMA posterior probabilities (**Figure 7C**) revealed strong support for four bilateral parameter sets. Top-down modulatory connections from STG to A1 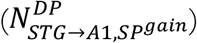 showed 96.07% posterior probability, indicating high confidence that systole modulates this connection. Amongst intrinsic gain parameters, SP gain in A1 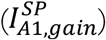 had 94.18% support, II gain in STG 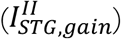 had 99.42% support, and SP gain in IFG 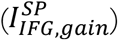 had 99.97% support. In contrast, other parameters showed weak support: top-down modulation from IFG to STG (36.34%), II gain in A1 (7.58%), SP gain in STG (47.49%), and II gain in IFG (33.18%).

The BMA posterior parameter estimates (**Figure 7D**) indicated the direction and magnitude of systolic-induced changes. The top-down modulatory connection 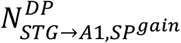 showed a negative effect, suggesting facilitation of self-inhibition that reduces the gain or precision of SP cells in A1 during systole. Positive effects were observed for 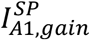 and 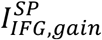, indicating increased self-inhibition in these regions during systole. A more pronounced positive effect was observed for 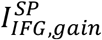, indicating a substantial increase in self-inhibition for the processing of tones coinciding with systole. Notably, there was also a marked increase in II gain at the level of STG 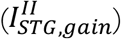, suggesting enhanced inhibitory control at this hierarchical level during systole. The effect on 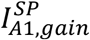 was modest and showed some hemispheric asymmetry (more pronounced in right A1), though we note the model was not configured to robustly infer hemispheric differences due to the prior specification of moderate hemispheric correlations.

These findings reveal a distributed pattern of gain control mechanisms underlying cardiac-specific effects. The strong support for both intrinsic gain parameters (particularly in A1 and IFG for SP cells, and in STG for II cells) and top-down modulatory connections indicates that systolic attenuation operates through multiple, complementary neural processes distributed across the auditory processing hierarchy. The predominance of intrinsic gain modulation (three parameters with >94% support) over extrinsic modulatory connections (one parameter with 96% support) suggests that cardiac-induced precision-weighting operates primarily through local cortical gain control mechanisms.

#### 3.2.3 Sensitivity Analysis

To understand how the model’s parameters captured cardiac-dependent effects, we performed sensitivity analysis on Model 10. This analysis examined how small perturbations to parameter values influenced the model’s predicted neuronal responses, quantifying each parameter’s contribution to systolic-induced attenuation.

##### Reconstructed responses

Figure 8 displays the model’s predicted responses for each cell population (SP, SS, II, DP) and the reconstructed local field potentials (LFPs) across the auditory hierarchy. The predicted LFPs showed cardiac-dependent effects most prominently at STG, with divergence between systolic and diastolic conditions emerging from 180 ms onwards. These effects were particularly evident in DP cell responses, suggesting that cardiac modulation substantially influences deep pyramidal dynamics at this hierarchical level.

**Figure 8.**
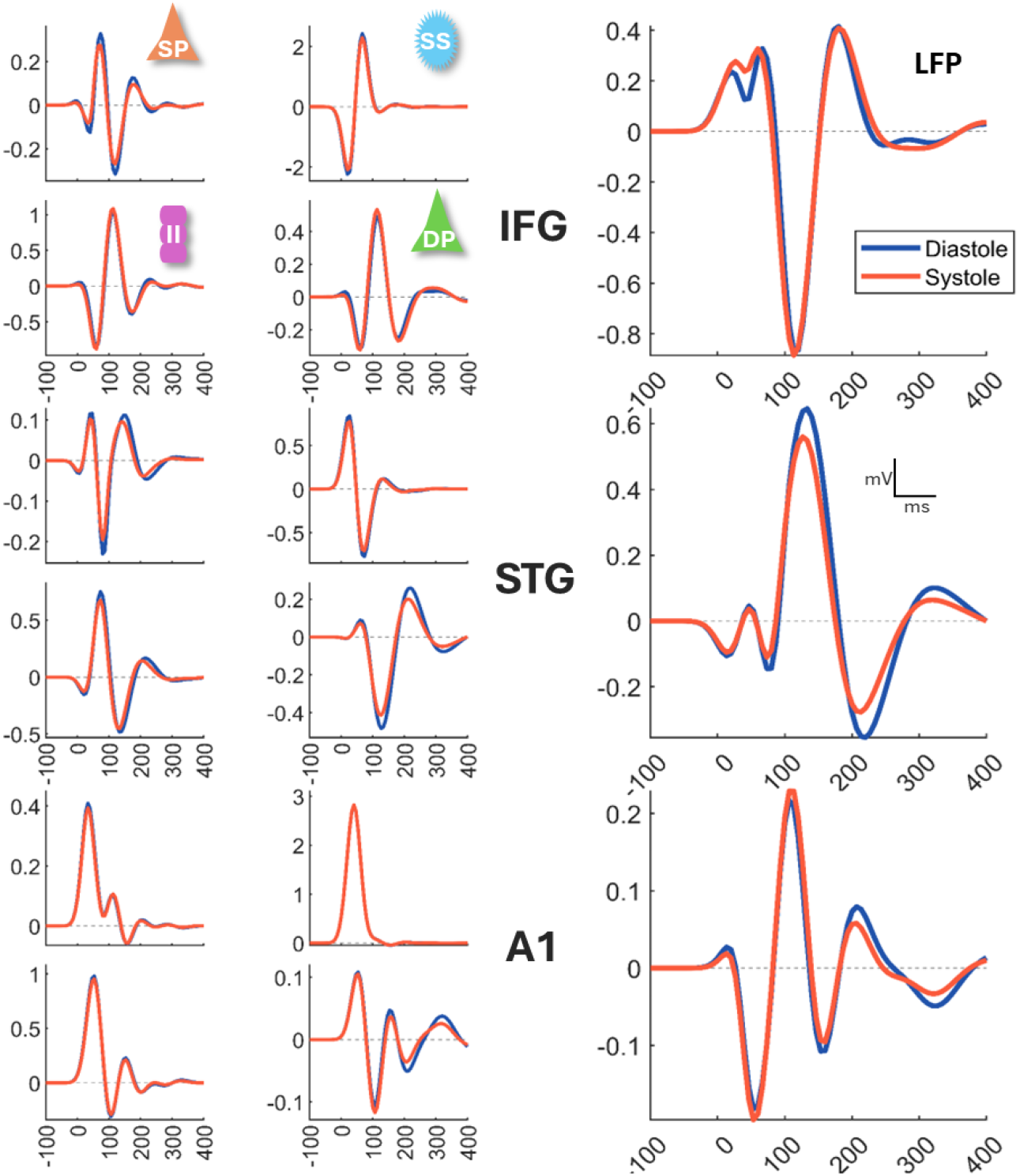
Reconstructed Peristimulus Responses: Model-predicted responses for primary auditory cortex (A1), superior temporal gyrus (STG), and inferior frontal gyrus (IFG). Left panels show responses for each cell population (SP, SS, II, DP) during systole (red) and diastole (blue). Right panels show reconstructed local field potentials (LFPs). Cardiac-dependent effects are most prominent at STG from 180 ms onwards.

**Figure 9.**
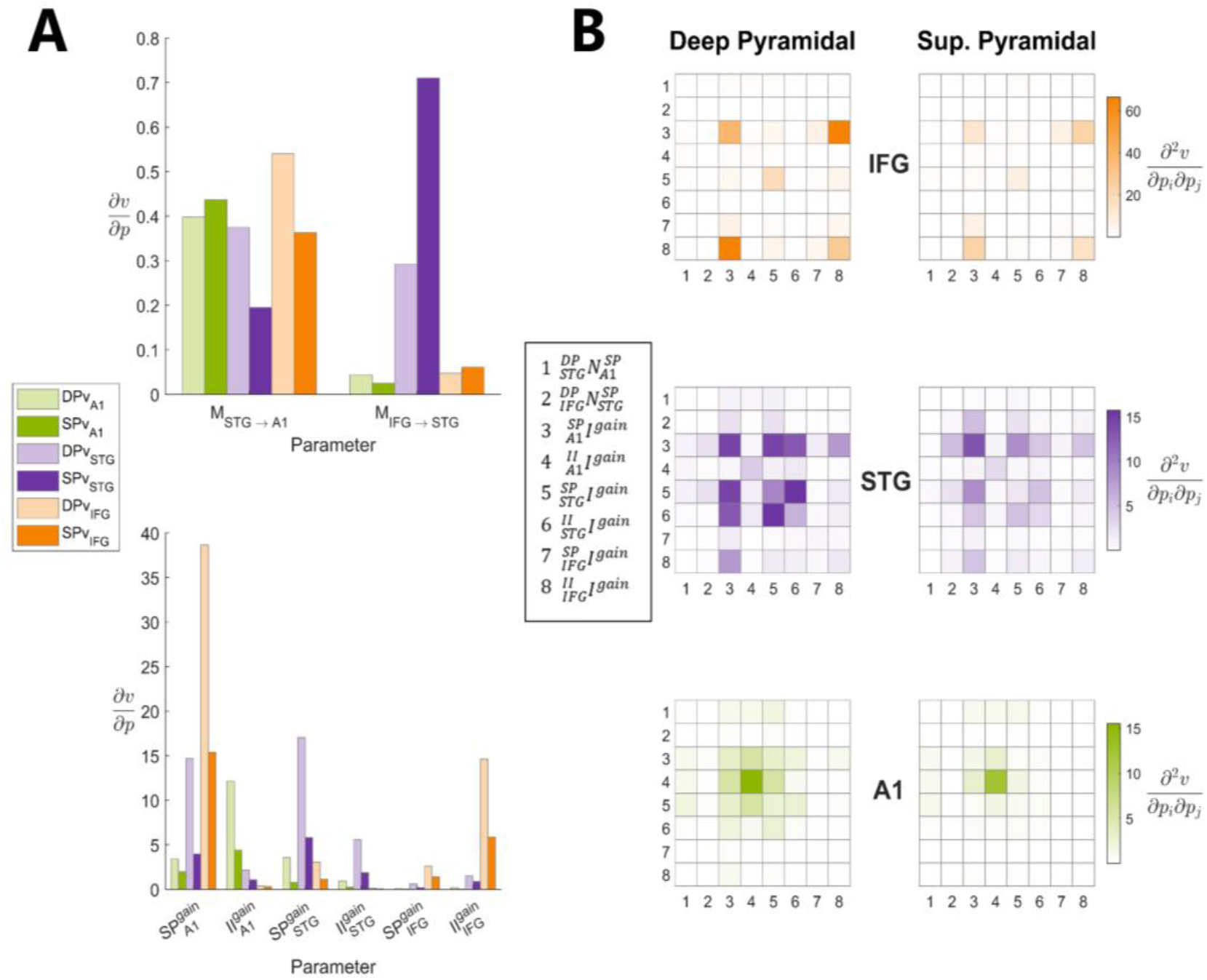
Sensitivity Analysis: (**A**) First-order sensitivity analysis showing direct effects of parameters on deep pyramidal (DP, upper panel) and superficial pyramidal (SP, lower panel) cell voltages. Intrinsic gain parameters (SP and II self-inhibition) show substantially larger effects than top-down modulatory connections. (**B**) Second-order sensitivity analysis showing parameter interactions at IFG (upper), STG (middle), and A1 (lower) for DP (left) and SP (right) cells. Colour intensity indicates interaction strength.

**Figure 9.**
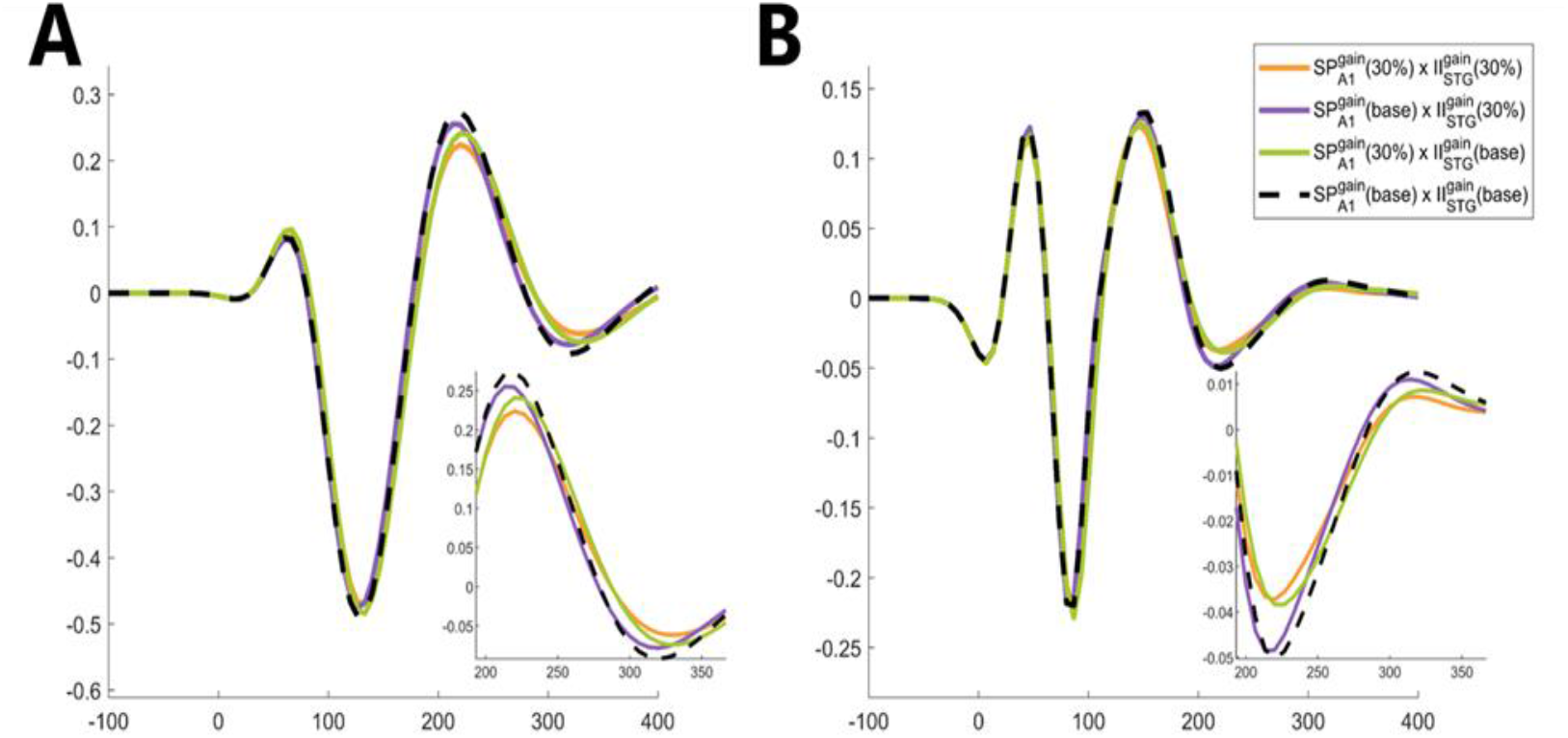
Parameter Contributions to Systolic Attenuation: Peristimulus responses at STG for (**A**) deep pyramidal and (**B**) superficial pyramidal cells under different parameter modulation conditions. Black dashed line shows baseline (no modulation). Coloured lines show effects of modulating 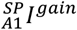 alone (green), 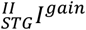 alone (purple), both parameters (orange), or neither (black). Insets show detail of late processing window (200-350 ms) where attenuation is most pronounced. Combined parameter modulation produces stronger attenuation than either parameter alone, demonstrating their synergistic interaction.

##### First-order sensitivity

We examined how individual parameters directly influenced DP and SP cell voltages by computing first-order partial derivatives (Figure A). This analysis revealed that intrinsic gain parameters had substantially larger effects than top-down modulatory connections. The contribution of modulatory connections (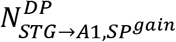 and 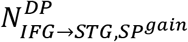) was an order of magnitude smaller than intrinsic gains. Amongst intrinsic parameters, 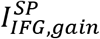 showed the largest direct effect, followed by 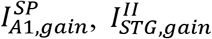, and 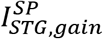. Notably, these effects were more pronounced on DP cells than SP cells across all hierarchical levels.

##### Second-order sensitivity

We examined parameter interactions by computing second-order partial derivatives (Figure B). Second-order interactions were sparse in A1 and IFG, whereas STG showed extensive interactions, indicating STG as a nexus to cardiac-dependent effects on auditory processing. Top-down modulatory connections showed minimal interactions with other parameters, including the notable absence of interaction between 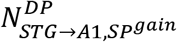 and 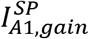. In contrast, intrinsic gain parameters exhibited substantial interactions across the hierarchy. 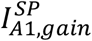 showed distributed influence, with its largest effect on DP cells in IFG and substantial interactions with 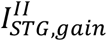 at the STG level. At the IFG level, interactions were most pronounced between 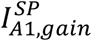 and 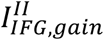, though the latter showed weak BMA support (33.18%), suggesting limited importance for cardiac-dependent effects.

##### Parameter contributions to systolic attenuation

To visualise how the key parameters identified by BMA contribute to systolic attenuation, we examined the peristimulus effects of modulating 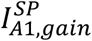 and 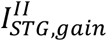 individually and in combination (Figure). Setting these parameters to 30% of their estimated systolic modulation revealed their functional impact on STG responses. 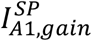 exerted substantial influence on STG dynamics, particularly during late processing (200-350 ms), producing marked attenuation in both DP and SP cells. 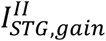 produced similar but smaller attenuation. Importantly, their combined modulation produced a stronger attenuating effect than either parameter alone, demonstrating that their interaction amplifies the systolic suppression of auditory processing at the STG level.

These sensitivity analyses demonstrate that systolic-induced attenuation operates primarily through intrinsic gain modulation rather than extrinsic modulatory connections. The distributed pattern of interactions, particularly between A1 SP gain and STG II gain, indicates that cardiac effects propagate through the auditory hierarchy via coordinated changes in local cortical excitability. This finding directly addresses our three candidate hypotheses from the Introduction: whilst top-down modulatory connections (NMDA-mediated mechanisms) contribute to cardiac modulation (96% posterior support), their functional impact is substantially smaller than intrinsic gain parameters. The dominance of intrinsic SP and II gain modulation, coupled with their synergistic interactions, provides strongest support for the inhibitory interneuron circuit hypothesis over extrinsic neuromodulatory mechanisms.

### 3.3 Summary

To summarise, sensor-space analyses revealed systolic-specific suppression of deviant responses, consistent with cardiac-phase-dependent precision-weighting of prediction errors. DCM model comparison identified an optimal architecture incorporating both top-down modulatory connections and intrinsic gain modulation in SP and II cells. Bayesian model reduction revealed that no single parameter configuration dominated (maximum posterior probability 19.8%), instead showing strong distributed support (posterior probabilities >94%) for SP gain in A1 (94%) and IFG (100%), II gain in STG (99%), and top-down modulation from STG to A1 (96%). Critically, sensitivity analysis demonstrated that intrinsic gain parameters exerted substantially larger effects than extrinsic modulatory connections, with SP gain in A1 and II gain in STG showing the strongest influences on network dynamics and exhibiting synergistic interactions. These findings indicate that cardiac-induced sensory attenuation operates through modulation of local cortical gain control mechanisms, with both intrinsic inhibitory interneuron circuits and top-down modulatory connections contributing to precision-weighting, though intrinsic mechanisms play the dominant role.

## 4 Discussion

We demonstrated that auditory responses to unexpected tones are selectively attenuated when they coincide with cardiac systole, specifically affecting the late mismatch negativity component at 200-250 milliseconds post-stimulus. Dynamic causal modelling revealed that this attenuation operates primarily through local cortical mechanisms: increased self-inhibition of superficial pyramidal cells, most prominent in the inferior frontal gyrus and primary auditory cortex, alongside enhanced inhibitory interneuron gain in the superior temporal gyrus. Top-down modulatory connections from STG to the gain of superficial pyramidal cells in primary auditory cortex also contributed, though sensitivity analysis showed that intrinsic gain mechanisms exerted substantially larger effects than top-down modulatory connections. This pattern suggests that cardiac-induced sensory attenuation operates primarily through local gain modulation rather than hierarchical precision-control. These findings provide the first empirical link between theoretical proposals about cardiac-sensory integration and specific synaptic mechanisms implementing precision-weighting in predictive processing.

### 4.1 Theoretical Interpretation: Precision-Weighting of Prediction Errors

The optimality of Bayesian belief updating depends on appropriately weighting prediction errors according to their reliability, formalised as precision-weighting (Feldman and Friston 2010; K. Friston 2005; Rao and Ballard 1999). The selective attenuation of deviant responses during systole, with no cardiac effects on standards, aligns with a precision-weighted predictive coding account of systolic-induced sensory attenuation. Standards generate minimal prediction errors regardless of reliability, whereas deviants generate substantial prediction errors that drive belief updating. If cardiac systole signals reduced sensory channel reliability, the brain should selectively downweight prediction errors generated during this period, whilst sparing predictable stimuli, precisely the pattern we observed.

The influence of cardiac state on late (200-250ms) exteroceptive sensory processing is in line with similar cardiac cycle studies (Al et al. 2020; Fouragnan et al. 2024; Kirasirova and Pyatin 2022; Ren et al. 2022). This finding indicates that systolic modulation affects post-perceptual synthesis rather than early sensory encoding. Late responses are typically associated with the updating of contextual representations (Halgren 1988; Pritchard 1981), attentional allocation (Polich 2007; Spencer and Polich 1999), arousal modulation (Conroy and Polich 2007) and belief updating in hierarchically higher-order regions (Garrido et al. 2009).

The delayed emergence of cardiac effects may reflect fundamental differences in how visceromotor versus skeletomotor states inform sensory processing, though we acknowledge this interpretation remains speculative. Saccadic eye movements provide a useful contrast: descending motor commands generate prospective predictions about upcoming sensory perturbations, enabling anticipatory attenuation of self-induced sensory consequences that modulate early sensory components (P1/N1, 80-200ms) (Perrinet, Adams, and Friston 2014; R. A. Adams et al. 2016; K. Friston and Kiebel 2009). Cardiac function, however, operates predominantly under involuntary autonomic control. While descending cardiovascular regulation exists via sympathetic and parasympathetic pathways, these slow-acting modulatory signals may not provide the precise millisecond-level predictions available for ballistic saccades.

This autonomic architecture potentially limits the brain’s ability to prospectively signal cardiac-induced sensory disturbances in the same manner as voluntary motor acts. Instead, the system may need to rely more heavily on retrospective estimation: inferring cardiac state and its sensory consequences from ascending interoceptive signals and the statistical properties of incoming prediction errors as they are generated, rather than from advance motor commands (K. Friston 2005; Parr and Friston 2019; Sridharan and Knudsen 2015). Under this tentative account, the brain would estimate current cardiac phase from available sensory evidence and then adjust precision-weighting accordingly, modulating the gain on prediction error signals based on inferred cardiac state. The late timing we observe would thus reflect when these retrospectively estimated precision weights are applied during hierarchical belief updating, rather than when cardiac state is initially detected.

However, several caveats warrant emphasis. First, our coarse-grained windowing approach (approximately 100ms windows) cannot definitively establish whether the 200-250ms timing reflects when cardiac state is inferred, when precision weights are computed, or simply when cardiac-modulated prediction errors become functionally significant for belief updating. The temporal smearing inherent in our analytical approach means earlier effects might be present but undetectable at our resolution. Second, this retrospective account does not preclude anticipatory components. Given the periodicity of the cardiac cycle and its ubiquity across the lifespan, the brain likely maintains generative models encoding temporal priors about cardiac dynamics. These learned priors could enable partial anticipation of cardiac-induced sensory fluctuations, filtering expected perturbations when a cardiac event is inferred based on recent history. Thus, cardiac-related sensory attenuation may reflect a hybrid inference process (partly retrospective, driven by real-time interoceptive evidence accumulation, and partly prospective, leveraging learned temporal structures of cardiac periodicity) that compensates for the absence of direct voluntary motor predictions through accumulated statistical knowledge of cardiac rhythms.

Disambiguating these mechanisms will require analytical approaches with finer temporal resolution and experimental manipulations that vary cardiac predictability. For instance, examining how heart rate variability affects the timing and magnitude of systolic attenuation could reveal whether the system relies on beat-to-beat inference or exploits longer-timescale regularities. Nevertheless, regardless of whether precision-weighting operates prospectively, retrospectively, or through hybrid mechanisms, our core finding remains: cardiac state systematically modulates the gain applied to prediction error signals during hierarchical auditory processing, consistent with precision-weighting accounts of interoceptive-exteroceptive integration.

### 4.2 Neurobiological Mechanisms

Our modelling indicated that deviant tones coinciding with systole are attributed reduced precision, evidenced by increased self-inhibition of SP cells in A1 and II cells in STG, with additional modulation from STG-to-A1 top-down connections. Sensitivity analysis revealed that intrinsic gains of SP cells in A1 were the primary drivers of systolic effects, with significant secondary contribution from II cells in STG, whilst top-down modulatory connections played only a minor role.

Within the canonical microcircuit framework (Bastos et al. 2012), superficial pyramidal cells encode prediction errors about causes from higher regions, spiny stellate cells encode prediction errors from lower regions, deep pyramidal cells compute expectations about hidden causes and states, and inhibitory interneurons encode prediction errors about hidden states. The attenuation of neural activity through increased self-inhibition of SP cells in A1 and increased inhibitory gain of II cells in STG aligns with this theoretical architecture. Specifically, reduced SP gain in A1 indicates decreased precision of prediction errors mapping causes between A1 and STG, whilst increased II gain in STG suggests this region contextualises cardiovascular influences on processing unexpected auditory stimuli. The identification of STG aligns with studies implicating this region in deviance detection (Blenkmann et al. 2019; Ishishita et al. 2019), attentional modulation of deviance detection (Sabri et al. 2006), and selective auditory attention (Neelon, Williams, and Garell 2006; Salmi et al. 2009). The specificity of STG for attenuating cardiophonic percepts is further corroborated by neuroimaging studies of pulsatile tinnitus showing aberrant superior temporal gyrus activity in response to cardiac-generated auditory signals (Lv et al. 2020; Xu et al. 2019).

Our modelling further indicates that response attenuation is most pronounced in DP cells of STG. As these cells represent expectations about causes and states, systolic-induced attenuation of DP activity reflects reduced belief updating about unexpected auditory events. This attenuation is mediated by increased II gain modulating deep-layer processes, which regulates local short-term plasticity as a function of salience (Jääskeläinen et al. 2007; Mongillo, Rumpel, and Loewenstein 2018). This mechanism aligns with recent DCM findings demonstrating that tonic inhibition regulates salient information processing with increasing prominence up the auditory hierarchy (N. E. Adams et al. 2020).

Previous DCM studies report that either NMDA or GABAergic transmission can encode expected information gains from exogenous contextual cues (Auksztulewicz and Friston 2015; Brown and Friston 2013). Our findings align more closely with a GABAergic account for endogenous contextual cues, as intrinsic gains influenced responses to a greater extent than top-down modulatory connections. This is consistent with positive associations between GABA concentration and interoceptive accuracy (Wiebking et al. 2014). A predominant GABAergic role may be expected given our passive auditory paradigm, where cardiovascular influences on auditory processing mirror normal experiences requiring no learning about this contingency that would necessitate NMDA-mediated long-term potentiation. Instead, our findings likely reflect a memory-based inferential process underlying systolic-specific effects that operates on the timescale of individual trials.

### 4.3 Implications and Clinical Relevance

Our findings bridge interoceptive inference and exteroceptive predictive processing, demonstrating that bodily states continuously modulate how external predictions are weighted. This has implications for understanding how physiological arousal affects perception and cognition across individuals.

Anxiety disorders frequently involve altered cardiac-sensory coupling, with anxious individuals showing heightened cardiac awareness and reduced ability to distinguish internal from external sensory events (Paulus and Yu 2012). Failed systolic attenuation may lead individuals to inappropriately update beliefs based on noisy signals during physiological arousal, contributing to hypervigilance and misattribution of internally-generated sensations to external threats. Similarly, somatic symptom disorders involve heightened attention to bodily sensations and difficulty differentiating internal from external events Wolters, Gerlach, and Pohl (2022)], potentially reflecting altered precision-weighting of interoceptive versus exteroceptive signals. Individual differences in intrinsic versus extrinsic gain mechanisms may correlate with interoceptive sensibility and clinical symptomatology, illuminating aetiological processes underlying these conditions.

Perhaps most directly translatable, pulsatile tinnitus, characterised by rhythmic sounds synchronised with heartbeat, may reflect failure of the attenuation mechanisms we identify. Our finding that STG implements cardiac-dependent gain control through inhibitory mechanisms suggests that dysfunction in these circuits could permit cardiac-generated acoustic and neuronal noise to reach perceptual awareness.

### 4.4 Limitations and Future Directions

Several limitations warrant consideration. We analysed grand-averaged event-related fields to maximise signal-to-noise ratio for fitting complex dynamic causal models, precluding assessment of individual differences. Future studies employing hierarchical dynamic causal modelling or parametric empirical Bayes could model individual variations and map interoceptive accuracy to synaptic mechanisms. Our coarse-grained windowing of cardiac phase based on electrophysiological markers assumed nearly instantaneous distributed effects throughout the body. In reality, different mechanisms operate with different latencies: baroreceptor activation peaks at 100-200ms post-contraction, cardiophonic effects at 200-300ms, and ballistic effects on brainstem at 300-500ms. Continuous convolution models or explicit generative models of cardiovascular physiology could disambiguate the relative contributions of these mechanisms.

We demonstrated neural attenuation but not behavioural consequences. Exploring performance in close-to-threshold detection and discrimination paradigms as a function of cardiac phase would test whether neural attenuation produces functionally significant perceptual effects. We focused on deviant responses based on sensor-level analyses showing cardiac-specific effects on deviants but not standards. Whilst theoretically motivated, this limits our ability to assess whether cardiac state modulates repetition suppression, which operates over multiple trials and might involve different synaptic mechanisms.

Future studies should examine whether cardiac phase modulates learning of statistical regularities, testing whether NMDA-mediated mechanisms implement multi-trial learning whilst GABA-mediated mechanisms handle within-trial perceptual inference adjustments. Introducing experimental conditions where statistical regularities are contingent on cardiac states would illuminate the biophysical mechanisms underpinning learnt relationships between visceral states and sensory processing. For instance, presenting tones either synchronously or asynchronously with heartbeats, or manipulating deviant probability contingent on cardiac phase, could test predictions about NMDA versus GABA mechanisms (Waselius et al. 2018). Such paradigms are particularly relevant for understanding hypervigilance to bodily sensations, a key factor impairing differentiation between external and internal sensory events in psychosomatic disorders (Wolters, Gerlach, and Pohl 2022).

Disambiguating prospective from retrospective cardiac-cued attention poses challenges due to difficulty experimentally controlling cardiac states. Leveraging generative models might facilitate this distinction: prospective attenuation could be influenced by recent cardiovascular events with earlier effects on sensory processing, whilst retrospective selection might depend on current cardiovascular state with minimal hysteresis and later effects on perceptual synthesis. This distinction could be validated by identifying discrepancies between model predictions and actual cardiac conditions, where prospective models provide efficient predictions with unsurprising cardiac events but are vulnerable to unexpected ones, and vice versa for retrospective models. Both modes likely operate concurrently, with their predominance possibly reflecting individual interoceptive capabilities. Manipulating cardiac predictability through biofeedback could further disambiguate these mechanisms. Clinical studies examining these processes in anxiety disorders, panic disorder, and somatic symptom disorders could illuminate how altered brain-body integration contributes to psychopathology and inform development of targeted interventions.

### 4.5 Conclusion

This study provides the first empirical characterisation of the synaptic and circuit-level mechanisms implementing cardiac-induced sensory attenuation within a predictive processing framework. We provide empirical evidence linking cardiac-induced sensory attenuation to specific cortical mechanisms implementing precision-weighting in predictive processing. Systolic attenuation of prediction error responses operates primarily through local inhibitory circuits in superior temporal gyrus rather than top-down hierarchical control, suggesting that bodily states directly modulate cortical gain in sensory hierarchies. These findings demonstrate that the body continuously shapes perceptual inference through mechanisms that operate largely outside awareness but may be disrupted in clinical conditions characterised by altered interoceptive-exteroceptive integration. Understanding these mechanisms at the level of synaptic processes and cortical circuits provides a foundation for developing interventions targeting disorders where brain-body interactions are compromised.

## Footnotes

1 Deviants per block were drawn from a non-homogeneous Poisson process. The rate parameter (*λ*) was sinusoidally modulated with a period equal to the block duration, varying between 3-11, with a minimum of two repetitions before deviants. This created clustering in portions of stimulus trains, with a randomised initial phase across blocks.

## Notes

### Competing Interest Statement

The authors have declared no competing interest.

